# The E3 ubiquitin ligase RNF216 contains a linear ubiquitin chain-determining-like domain that functions to regulate dendritic arborization and dendritic spine type in hippocampal neurons

**DOI:** 10.1101/2023.10.19.563080

**Authors:** Jayashree Kadirvelu, Savannah E. Jacobs, Ruochuan Liu, Antoinette J. Charles, Jun Yin, Angela M. Mabb

## Abstract

Of the hundreds of E3 ligases found in the human genome, the RING-between RING (RBR) E3 ligase in the LUBAC (linear ubiquitin chain assembly complex) complex HOIP (HOIL-1-interacting protein or RNF31), contains a unique domain called LDD (linear ubiquitin chain determining domain). HOIP is the only E3 ligase known to form linear ubiquitin chains, which regulate inflammatory responses and cell death via activation of the NF-κB pathway. We identified an amino acid sequence within the RNF216 E3 ligase that shares homology to the LDD domain found in HOIP (R2-C). Here, we show that the R2-C domain of RNF216 promotes self-assembly of all ubiquitin chains, with a dominance for those assembled via K63-linkages. Deletion of the R2-C domain altered RNF216 localization, reduced dendritic complexity and changed the distribution of apical dendritic spine morphology types in primary hippocampal neurons. These changes were independent of the RNF216 RBR catalytic activity as expression of a catalytic inactive version of RNF216 had no effect. These data show that the R2-C domain of RNF216 diverges in ubiquitin assembly function from the LDD of HOIP and and functions independently of RNF216 catalytic activity to regulate dendrite development in neurons.

## Introduction

Ubiquitination is a posttranslational modification that involves the covalent attachment of the 76-amino acid protein called ubiquitin to other protein targets (1). Protein ubiquitination regulates a broad range of cellular processes and is essential for numerous aspects of nervous system development and regulation (2, 3). In most cases, protein ubiquitination relies on an enzymatic series cascade that includes an ubiquitin activating enzyme (E1), ubiquitin conjugating enzymes (E2s) and ubiquitin ligases (E3s) (1). The complementation of this cascade is responsible to drive substrate specificity, which is strongly dictated by the E3. Moreover, varying combinations of the ubiquitin enzyme cascade creates a diversity of assembled ubiquitin moieties on their substrates thus creating a broad range of cellular signaling events (4). While the quantity of genes that encode for E1 and E2s is relatively small (∼40), the number of genes encoding E3s is greater than 10-fold higher (∼600-800) (5). Importantly, E3s have a diversity of catalytic domains that include CRL, DDB-Box, SOCS Box, HECT, RING and RBR (6).

In addition to identifying substrates for the massive number of E3s, other limitations in the ubiquitin field relate to determining how their catalytic domains function to ubiquitinate their targets and how these domains can generate a diversity of ubiquitin linkages. For example, the RBR E3, RNF216/TRIAD3 has recently been demonstrated to assemble K11-, K48-, and K63-linked ubiquitin chains onto its targets (7–9), which may be controlled in part by its phosphorylation (7). Here, we attempted to delineate the function of the RNF216 RBR catalytic region in regulating its ubiquitination. In our analysis, we identified an amino acid sequence juxtaposed to the RNF216 RBR domain (R2-C) that shares homology to a linear ubiquitin chain determining domain (LDD) domain found in the RBR E3 HOIP (HOIL-1-interacting protein or RNF31), which is a component of the LUBAC (linear ubiquitin chain assembly complex) complex. HOIP is the only E3 known to form linear ubiquitin chains, which regulate inflammatory responses and cell death via activation of the NF-κB pathway (10, 11). Here, we evaluated a possible role for the R2-C of RNF216 in assembling linear ubiquitin chains.

However, instead, we found that the R2-C domain promotes self-assembly of all ubiquitin chains, with a dominance for those assembled via K63-linkages. Deletion of the R2-C domain altered RNF216 localization in primary hippocampal neurons, reduced dendritic complexity and changed the distribution of apical dendritic spine morphology types in primary hippocampal neurons. These changes were independent of the RNF216 RBR domain activity, as expression of a catalytic inactive version of RNF216 had no effect. These data show that the R2-C domain of RNF216 diverges in ubiquitin assembly function from the LDD of HOIP and functions independently of RNF216 catalytic activity to regulate dendrite development in neurons.

## Materials and Methods

### Antibodies and Reagents

Chemicals and reagents were purchased from Sigma-Aldrich, Thermo Fisher Scientific, Invitrogen, or Santa Cruz Biotechnology. Cell culture reagents were obtained from Gibco^TM^ (Thermo Fisher Scientific), unless otherwise stated. Anti-ubiquitin (P4D1) (Santa Cruz Biotechnology, sc-8017), anti-Flag (Sigma-Aldrich, F7425; Sigma-Aldrich, F1804), HA.11 epitope tag (Biolegend, 902301), anti-ubiquitin (Novus Biologicals, NBP2-30132), c-myc (SC Biotech, SC-40; Bethyl, A190-104A), and Beta Actin (GT5512) (GeneTex, GTX629630) were used as primary antibodies. Secondary antibodies from LI-COR Biosciences include donkey anti-goat IgG (Li-COR, 926-68074), donkey anti-rabbit IgG (Li-COR, 925-68073), goat anti-mouse IgG (Li-COR, 926-68070), goat anti-rabbit IgG (Li-COR, 926-32211), and donkey anti-mouse IgG (Li-COR, 926-32212). Secondary antibodies for immunostaining were Goat anti-Mouse IgG1 Secondary Antibody, Alexa Fluor 647 (Invitrogen, A21240) and DAPI (Thermo Fisher Scientific, 62248).

### Plasmid Construction

The original Uba1 plasmid was a gift from Dr. Wade Harper of Harvard Medical School (12) and the original Ubch7 and HA-UB plasmids were kind from Dr. Bernard Roizman and Suzanne Cohen, respectively, of the University of Chicago (13, 14). The E6AP plasmid was a gift from Dr. Jon M. Huibregtse of the University of Texas at Austin (15). The pET28a-wtUba1, pET21b-wtUbcH7, and pET28a-HA-UB constructs were cloned as previously described (16). The pET28a-flag-E6AP HECT domain construct was cloned as previously described (17).

A FLAG tag sequence (DYKDDDDK) was added to the N-terminus of hRNF216 in the pOPINB-hRNF216 construct, a gift from Dr. Bernard Lechtenberg (Addgene plasmid # 167354; http://n2t.net/addgene:167354; RRID:Addgene_167354) (7) via amplification of the hRNF216 gene with forward primer RL1, encoding the FLAG and 6XHis tag sequences, and reverse primer RL2, using a Polymerase Chain Reaction (PCR). The PCR product was digested with enzymes NcoI and HindIII (New England Biolabs, Ipswich, Massachusetts, USA), ligated back into the pOPINB vector with T4 DNA ligase (New England Biolabs), transformed into XL1-Blue cells (Agilent Technologies, Santa Clara, California, USA) followed by DNA extraction using a Qiagen plasmid mini kit (Qiagen, Crawley, UK), and the novel construct confirmed through Sanger Sequencing via Azenta/GENEWIZ (Azenta Life Sciences Inc., South Plainfield, New Jersey, USA).

For the hRNF216 truncations, all were amplified with the same forward primer RL3, encoding an N-terminal FLAG and 6XHis tag sequence. While the forward primer for PCR amplification was identical for all the truncations, each was constructed with its own unique reverse primer, all of which culminated with a stop codon after the indicated C-terminal residue of the truncate: the plasmid hRNF216 (501–722) was amplified with reverse primer RL4, hRNF216 (501–736) was amplified with reverse primer RL5, and hRNF216 (501–866) was amplified with reverse primer RL2. All three PCR product inserts of the truncates were digested utilizing the same restriction enzymes, NcoI and HindIII (New England Biolabs), and each truncation was ligated back into the pOPINB vector, transformed into XL1-Blue cells (Agilent).

The pET22b-UB K48R and pET22b-UB K63R were kind gifts from Dr. Eric Strieter of the University of Massachusetts at Amherst (18). The pET28a-HA-UB construct which was cloned as previously described (16). The ubiquitin K to R mutants we generated in the same vector as the wtUB construct, with the HA tag sequence in the same position at the UB N-terminus to ensure the protein purification and subsequent *in vitro* ubiquitination assays would be performed in the same manner and could be immunoblotted with an HA or ubiquitin specific antibody. The UB K48R mutant and UB K63R mutant were PCR amplified using the same primers: forward primer SJ1 encoding the HA tag, and reverse primer SJ2. Both UB K48R and UB K63R PCR inserts were digested with Nde1-HF and NotI-HF (New England Biolabs), ligated into the pET28a vector, transformed into XL1-Blue competent cells (Agilent).

pRK5-FLAG-RNF216 was a gift from Dr. Tsung-Hsien Chuang (Sanford-Burnham Medical Research Institute). To make FLAG-tagged RNF216 mutants for expression in mammalian cells, the pRK5-FLAG-RNF216 construct was cut with SalI and BamHI restriction enzymes to isolate the pRK5-FLAG vector. The inserts were PCR amplified and then subcloned into the pRK5-FLAG vector to generate ΔR2-C, RBR-R2-C, and RBR only. Plasmids were validated by DNA sequencing. For all generated plasmids, DNA was extracted using a Qiagen plasmid mini kit (Qiagen). Confirmation of the correct constructs was performed with Sanger Sequencing via Azenta/GENEWIZ (Azenta Life Sciences Inc.).

### Protein purification

Each plasmid was transformed into BL21 (DE3)pLysS chemically competent *Escherichia Coli* cells (Invitrogen, La Jolla, California, USA Cat# C606010) via electroporation with recovery in SOC (Research Products International, Mt. Prospect, Illinois, USA) followed by incubation at 37°C for one hour with agitation, and subsequently plated on LB-agar plates (Research Products International) with the appropriate antibiotic selection marker.. A starting culture was prepared using 2XYT Broth media (Research Products International) with a 1:1000 dilution of the appropriate antibiotic and a single colony from the relevant LB Agar plate and incubated at 37°C overnight with agitation. The following day, a 1:1000 dilution of antibiotic was added to 1 L of 2XYT broth media in addition to a 1:200 dilution of starter culture. The 1 L culture was incubated at 37°C with agitation for several hours until reaching mid-log phase, (an OD_600_ of approximately 0.6-0.8) at which point isopropyl ß-D-1-thiogalactopyranoside (IPTG) was added for induction to a final concentration of 1 mM, the temperature decreased to 25°C, and the culture allowed to grow overnight with agitation. The following day, the culture was spun down for 20 minutes at 5,500 rpm, and the supernatant discarded. Ni-NTA lysis buffer was added to each centrifuge bottle to fully resuspend each cell pellet and the contents of each bottle combined. Then, lysozyme (1 mg/mL of cell lysate), 1 mM PMSF, and cOmplete EDTA-free Protease Inhibitor Cocktail, mini-Tablet (Sigma-Aldrich, Burlington, Massachusetts, USA), followed by brief vortexing, and incubation on ice for 30-60 minutes. After incubation on ice, the lysate was sonicated at 30% amplitude with alternating 10 second pulses for 15-30 minutes or until the color of the lysate darkened. The lysate was then centrifuged at 12,000 rpm for 60 minutes, the supernatant decanted into a fresh tube and the pellet discarded. The protein was then purified on Ni-NTA affinity chromatography with Ni-NTA agarose resin (Qiagen) following the protocol described in *Qiagen*, 2003, and dialyzed overnight. After the protein was collected the following day and concentrated using 10 kDa MWCO Amicon Ultra (Merck Millipore). The protein concentration was monitored periodically during the process using Bio-Rad Protein Assay Dye Reagent Concentrate (Bio-Rad, Hercules, California, USA) and concentrated to approximately 100-200 μM.

The RNF216 recombinant proteins were expressed in the same manner with some modifications, as described in *Cotton et al*., 2022 (7) with the following modifications: First, once the 1 L cell culture reached an OD_600_ of ∼0.8, the temperature was decreased to 20°C for 45 minutes prior to the addition of 0.5 mM ZnCl_2_ and 0.5 mM IPTG. Second, as this particular protein (and even its smaller RBR truncations) are prone to aggregation and inclusion body formation, the composition of all the Ni-NTA buffers (including the lysis, wash, and elution buffers) was slightly modified by the addition of 10% glycerol and 10% sucrose. While some proteins may have enhanced yield by binding to Ni-NTA resin for a few hours or even overnight, RNF216’s large size and penchant for aggregation contraindicate extended binding. Therefore, we performed the Ni-NTA affinity column chromatography immediately after sonication and centrifugation. Finally, an additional 1 mM of dithiothreitol (DTT) was added to the final protein product for storage at –80°C to aid in maintaining and prolonging the activity of recombinant RNF216.

## *In vitro* ubiquitination assays

### Control autoubiquitination assays

Control reactions were performed to ensure that the purified recombinant proteins had adequate activity. Because RNF216 is believed to have a bias for the formation of K63-polyubiquitin chain linkages, the HECT domain of the E3 ubiquitin ligase UBE3A was used in all assays to serve as a comparison against RNF216 and its truncations as UBE3A has a well-documented propensity for the formation of K48 polyubiquitin chains linkage types.

### Non-stringent control assays

We first tested the proteins under less stringent conditions to ensure their activity. These reactions were performed by combining 1X Tris-HCL/MgCl_2_ reaction buffer, 5 mM ATP, 1mM DTT, followed by the addition of 1 μM recombinant Uba1, 3 μM recombinant UbcH7, 5 μM of each E3, including recombinant flag-E6AP HECT domain, recombinant flag-RNF216 FL, recombinant flag-RNF216 (501–722), recombinant flag-RNF216 (501–736), and recombinant flag-RNF216 (501–866), and finally, 50 μM of HA-wtUB. Sterile cell culture H_2_O was added to a total reaction volume of 50 μL for each reaction. For each E3 ligase, additional control reactions were utilized for comparison in which one of the four protein components was eliminated: the first reaction for each E3 contained all components, the second reaction eliminated Uba1, the third reaction eliminated UbcH7, and the fourth reaction eliminated the E3 ligase. After all components were added to each reaction and adequately mixed, the reactions were incubated at 37°C with agitation for four hours. They were subsequently quenched through the addition of 6x DTT loading dye and boiled for a minimum of five minutes. SDS-PAGE was performed for each reaction followed by analysis of the immunoblots probed with anti-HA antibody (Santa Cruz Biotechnologies, Dallas, Texas, USA). If all proteins are active, the reaction with all components should result in the appearance of a very strong smear while elimination of Uba1 should result in no smear as there is no enzyme present containing an adenylation domain to activate ubiquitin with ATP for subsequent transthiolation to the UbcH7 active site. The reaction eliminating UbcH7 may show a weak smear as, unlike ubiquitination cascades involving RING or Ubox E3 ligases in which E2-conjugating enzymes play a crucial role, many HECT and RBR E3’s are capable of capturing ubiquitin directly from an E1 enzyme in the absence of an E2 enzyme (albeit, much less effectively) due to the presence of a catalytic cysteine in HECT and RBR E3 ligases. Elimination of the E3 ligase will also show little to no smear as most E2-conjugating enzymes are only capable of being charged with a single ubiquitin molecule covalently bound to the catalytic cysteine residue in their active site. However, many E2s are also capable of binding a second ubiquitin noncovalently to a ß-sheet on their “backside” or, essentially, on the opposite face of the E2 distal from the active site. The purpose of this second, noncovalently bound ubiquitin has yet to be fully elucidated, but is theorized to endow E2s with the ability to processively transfer polyubiquitin chains directly onto a RING/Ubox E3 for autoubiquitination or directly onto a substrate of the RING/Ubox E3 ligase. Due to the potential for noncovalent “backside” binding ubiquitin on some E2’s in addition to a charged E2’s covalently bound ubiquitin, the appearance of a di-UB or, if E1 is also loaded, tri-UB band in reactions without the E3 component is not uncommon.

### Stringent control assays

The stringent control assays were performed in the same manner as the non-stringent control assays with the only difference being the concentrations of the four protein components and incubation time. These reactions were performed by combining 1X Tris-HCL/MgCl_2_ reaction buffer, 5 mM ATP, and 1mM DTT; however, for this set of reactions, only 0.1 μM recombinant Uba1, 0.5 μM recombinant UbcH7, 2 μM of each E3, including recombinant flag-E6AP HECT domain, recombinant flag-RNF216 FL, recombinant flag-RNF216 (501–722), recombinant flag-RNF216 (501–736), and recombinant flag-RNF216 (501–866), and 3 μM of HA-wtUB were added. As in the above control reactions, sterile cell culture H_2_O was added to a total reaction volume of 50 μL for each reaction. After all components were added to each reaction and adequately mixed, the reactions were incubated at 37°C with agitation; but only for 90 minutes instead of four hours. The reactions were subsequently quenched by the addition of 6x DTT loading dye and boiled for a minimum of five minutes. SDS-PAGE was then performed for each reaction followed by analysis of the immunoblots probed with anti-HA antibody (Santa Cruz Biotechnologies). The same control reactions described above were also used for these reactions.

### In vitro autoubiquitination assays using chain linkage specific antibodies

After having verified that all proteins were active, we proceeded to examine the chain-linkage specificity of the FL RNF216 and if or how our various truncations of the RNF216 RBR domain might affect this chain specificity. The UBE3A HECT domain, which forms K48 polyubiquitin chains, was used to compare. In this set of reactions, linkage specificity was analyzed through the utilization of K48 and K63 polyubiquitin linkage specific antibodies. We used the non-stringent set of conditions for these assays. These reactions were performed by combining 1X Tris-HCL/MgCl_2_ reaction buffer, 5 mM ATP, 1mM DTT, followed by the addition of 1 μM recombinant Uba1, 3 μM recombinant UbcH7, 5 μM of each E3, including recombinant flag-E6AP HECT domain, recombinant flag-RNF216 FL, recombinant flag-RNF216 (501–722), recombinant flag-RNF216 (501–736), and recombinant flag-RNF216 (501–866), and finally 50 μM of HA-wtUB. Sterile cell culture H_2_O was added to a total reaction volume of 50 μL for each reaction. After all components were added to each reaction and adequately mixed, they were incubated at 37°C with agitation for four hours. The reactions were subsequently quenched by as stated above and loaded onto three separate SDS-PAGE gels. Immunobloting was performed, probing one with anti-HA antibody (Santa Cruz Biotechnologies), one with anti-K48-linkage Specific Polyubiquitin (D9D5) Rabbit (Cell Signaling Technologies, Danvers, MA, USA), and third with K63-linkage Specific Polyubiquitin (D7A11) Rabbit (Cell Signaling Technologies).

### In vitro autoubiquitination assays using ubiquitin K to R mutants

Like the assays using the linkage specific antibodies, it is necessary to run all the reactions with each E3 on the same blot to compare activity, so only the reactions with all components and the reactions eliminating Uba1 were used. As described previously, these reactions were performed by combining 1X Tris-HCL/MgCl_2_ reaction buffer, 5 mM ATP, and 1mM DTT, followed by the addition of 0.1 μM recombinant Uba1, 0.5 μM recombinant UbcH7, 2 μM of each E3, including recombinant flag-E6AP HECT domain, recombinant flag-RNF216 FL, recombinant flag-RNF216 (501–722), recombinant flag-RNF216 (501–736), and recombinant flag-RNF216 (501–866). Reactions were performed in triplicate. To each of the three identical triplicates, 3 μM of one UB construct were added, including HA-wtUB to serve as a control, as well as HA-UB K48R and HA-UB K63R. Sterile cell culture H_2_O was added to a total reaction volume of 50 μL for each reaction. After all components were added to each reaction and adequately mixed, the reactions were incubated at 37°C with agitation for 90 minutes. The reactions were subsequently quenched as stated above. Three separate SDS-PAGE gels were run, with one gel harboring all of the reactions with HA-wtUB, a second with all the reactions using the HA-UB K48R mutant, and the third with the reactions using the HA-UB K63R mutant. After SDS-PAGE, each immunoblot was probed with anti-HA antibody (Santa Cruz Biotechnologies) and analyzed.

### Cell culture (Primary Hippocampal Neurons and HEK293T Cells)

Primary hippocampal neurons were cultured as described as previously described (Wall et al 2018). HEK293T cells were cultured in DMEM with 10% fetal bovine serum and 1% Pen Strep at 37 °C.

### Transfection and Preparation of Cell Cultures

#### Primary Hippocampal Neurons

Primary hippocampal neurons from *Rnf216/Triad3^fl/fl^*postnatal day 0 and 1 mouse pups were prepared as previously described (Wall et al, 2018). For dendritic complexity and dendritic spine studies, neurons were transfected at DIV13 using Lipofectamine 2000 (Thermo Fisher Scientific) as described previously (Wall et al 2018). Neurons were transfected with tdTomato-P2A-CRE, tdTomato-P2A-CRE/Flag-RNF216 WT (wildtype), tdTomato-P2A-CRE/Flag-RNF216 CA (catalytically inactive) (19), and tdTomato-P2A-CRE/Flag-RNF216 ΔR2-C. At DIV17, neurons were fixed with 4% PFA/4% Sucrose.

#### HEK293T Cells

0.25 x 10^6^ HEK293T cells were transferred to each well of a 6-well dish containing 2mL DMEM with 10% fetal bovine serum and 1% Pen Strep. The cells were then incubated at 37 °C for 24 hours. After 24 hours, cells were transfected using Lipofectamine 3000 and P3000 transfection reagents. Cells were harvested with ice cold DPBS and a cell scraper after incubation at 37 °C for 24 hours.

### Protein Extraction and Western Blotting

#### For in vitro ubiquitination assays

For SDS-PAGE and subsequent western blot analysis, equal amounts of reaction product were loaded in each lane of a 4-15% SDS-PAGE gel (Bio-Rad). The gels were run in a 1X Tris/glycine/SDS buffer, followed by a semi-dry transfer of the proteins onto a polyvinylidene fluoride (PVDF) membrane (Bio-Rad) using Trans-Blot Turbo Transfer System (Bio-Rad).

Following transfer, the PVDF membrane was blocked for one-hour in a 5% non-fat dry milk solution made with TBST. The membrane was then incubated overnight at 4°C with agitation in a 5% non-fat dry milk TBST solution with the addition of 1:1000 dilutions of each of the primary antibodies employed in this study. Following overnight incubation, the membranes were washed with TBST three times for five minutes each, and subsequently incubated for one-hour at room temperature with agitation in a 5% non-fat dry milk TBST solution containing a 1:20,000 dilution of either goat anti-mouse IgG-HRP for immunoblots using anti-HA as the primary antibody or goat anti-rabbit IgG-HRP for immunoblots using either of the polyubiquitin chain linkage specific antibodies. After incubation with the secondary antibody, membranes were washed with TBST again three times for five minutes each followed by chemiluminescent detection with SuperSignal® West Pico Chemiluminescent Substrate (Thermo Fisher Scientific, Waltham, Massachusetts) on a ChemiDoc Touch Imaging System (Bio-Rad)

#### For protein expression in HEK293T cells

Transfected HEK293 cells were lysed with RIPA Buffer (50mM Tris HCl, 150mM NaCl, 0.15% SDS, 1% NP40, 0.5% sodium deoxycholate, pH 8.0) and cell extracts were centrifuged at 14,000 rpm for 10 min at 4°C. Supernatants were collected and protein concentration was measured using Pierce 660 nm protein assay kit (Thermo Fisher Scientific). 2X SDS Loading Buffer was added to equal amounts of protein samples and analyzed by running on a 4-15% 10-well SDS-PAGE gel (Bio-Rad). Gels were run for and then transferred for 1 hour at 70mV. The nitrocellulose membrane was blocked overnight with Li-COR blocking buffer. Primary antibodies were added and incubated overnight at 4°C. Secondary antibodies and incubation were conducted as previously described (20).

### Immunoprecipitation

#### Immunoprecipitating Myc-Arc

pcDNA3.1, Flag-Triad3A, HA-Ubiquitin, myc-Arc, and the flag-tagged plasmid constructs were transfected in HEK293 cells by Lipofectamine 3000 (Invitrogen). IP buffer (20 mM Tris-HCl, 3 mM EDTA, 3 mM EGTA, 150 mM NaCl, 1% Triton X-100, pH 7.4) with 2 mM DTT, 1μg aprotinin/mL, 1:2000 of 10mg/mL stock of leupeptin, 0.5 mM PMSF, 2-5mM N-ethylmaleimide (NEM), 2-3mM Iodoacetamide (IAA), and 2000 Units/mL DNase was prepared. Cells were harvested 24 hours after transfection and lysed for 20 minutes at 95°C on a heat block in ubiquitin IP buffer consisting of stock IP buffer described above and 1% SDS. Samples were cooled and then spun down at 13,000 rpm for 20 minutes. The supernatants were taken and then diluted 10 fold in IP buffer. Anti-myc antibody was added to the samples and allowed to tumble for 1 hour at 4°C. Gamma Bind G-Sepharose beads (GE Healthcare Life Sciences) were then added and samples were tumbled overnight at 4°C. After washing with IP buffer three times, immunoprecipitants were boiled in 2X SDS sample buffer in a 1:1 ratio, separated by SDS-PAGE electrophoresis, transferred to a nitrocellulose membrane, and immunoblotted with anti-myc (Abcam), anti-FLAG (Sigma), or anti-ubiquitin antibodies.

#### Immunoprecipitating flag-tagged constructs

HEK293 cells were transfected with pcDNA3.1, FLAG-Triad3A, and flag-tagged plasmid constructs with the procedure described above. After denaturing immunoprecipitation using the above protocol, the products were boiled in 2X SDS sample buffer in a 1:1 ratio, separated by SDS-PAGE electrophoresis, transferred to a nitrocellulose membrane, and immunoblotted with anti-FLAG (Sigma), and anti-Ub P4D1 antibodies.

### Immunocytochemistry

#### Cells for colocalization of flag-tagged protein

Transfected hippocampal neurons were fixed at DIV17 for 20 minutes in ice cold DBPS containing 4% paraformaldeyde/4% sucrose in 1XPBS at 4°C. For immunostaining, cells were permeabilized with 0.2% saponin in PBS for 15 minutes at room temperature and then washed three times with DPBS. Neurons were blocked in 10% NGS for 1 hour at 37°C. Neurons were incubated with mouse anti-FLAG (Sigma) antibody overnight at 4°C. Cells were washed three times with DPBS and incubated with Goat anti-Mouse IgG1 Alexa Fluor 647 Secondary Antibody and DAPI (Thermo Fisher Scientific) for 1 hour at room temperature. Following incubation, neurons were washed 3 times with 3% NGS and then mounted on slides as previously described (20).

#### Cells for dendritic spine morphology

Cultured hippocampal neurons were prepared as previously described. Cells were washed three times with DPBS and incubated with Goat anti-Mouse IgG1 Secondary Antibody, Alexa Fluor 647 (Invitrogen) and DAPI (Thermo Fisher Scientific) for 1 hour at room temperature.

### Image Acquisition and Analysis

Western blots were analyzed with Fiji Is Just ImageJ (FIJI). For dendrites, images of fixed samples of primary hippocampal neurons were acquired with an LSM 700 (Zeiss) confocal microscope using 20X objective and analyzed using FIJI. For dendritic spines, images of fixed samples of primary hippocampal neurons were acquired with an LSM 700 (Zeiss) confocal microscope using 63X objective. Images for dendritic spines were analyzed using FIJI and Imaris 9.2. Specifically, only the primary branches of apical or basal dendrites were analyzed. Spine density was calculated by number of spines per unit length of dendrite. Spine volume and proportion of each spine type was also calculated with numerical data provided by Imaris 9.2.

### Statistical Analysis

All numerical data were expressed as a mean with SEM. Statistical analyses were conducted with GraphPad Prism8 software (GraphPad Software, San Diego, CA). Comparison between groups were assessed with ANOVA, followed by Tukey’s post hoc test. In all graphs, significance was set at p < 0.05.

## Results

### RNF216 contains an LDD-like domain that shares homology to the LDD domain of HOIP

HOIP is a member of the RBR E3 family (Figure 1A) and is the only known E3 ligase to assemble linear ubiquitin chains via its LDD domain (10, 11). Thus, we reasoned that perhaps other RBR E3 ligases may also assemble linear ubiquitin chains. Upon conducting a systematic search, we identified a region lying adjacent to the RBR domain of the E3 ligase RNF216 with shared homology to the LDD domain of HOIP (Figure 1B). LDD domain homology included the RBR-C-helix of RNF216 (Figures 1C and 1D) (7, 21).

**Figure 1:**
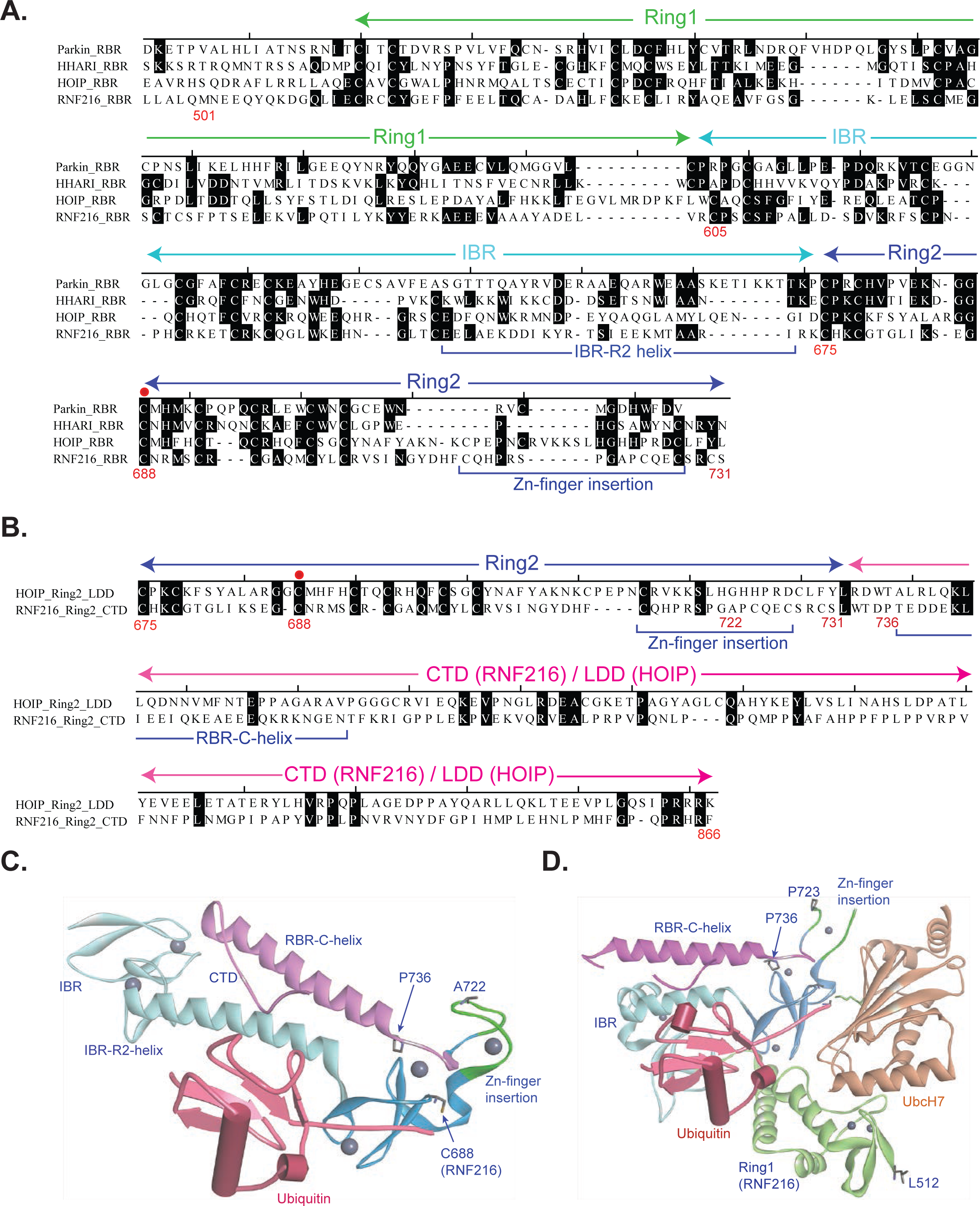
Identification of a putative LDD-like domain in RNF216. (A) Alignment of RBR E3s that include Parkin, HHARI, HOIP, and RNF216. (B) Alignment of the LDD domain of HOIP with the C-terminal region of RNF216. RNF216 shares some degree of homology to the LDD domain of HOIP, which includes the RBR-C-helix (R2-C) of RNF216. (C) Modified crystal structure from Cotton et al (PDB 7M4M) (7) showing the position of the RBR-C-helix (R2-C, magenta) domain in relation to the Zn^2+^ finger regions and Ubiquitin (UB, fuchsia). The R2-C domain may help coordinate Zn^2+^ within the Ring 2 domain and may help to facilitate the binding of UB to the IBR-R2 α-helix. Zn^2+^ ions are shown as gray balls. (D) Modified crystal structure from Cotton et al (PDB 8EBO) (21) showing the position of the R2-C domain when RNF216 is bound to its E2 (UbcH7, beige).

To determine if this C-terminal LDD-like domain alters RNF216 function, we generated a series of RNF216 mutants (Figure 2A). Using an in vitro ubiquitin assay with purified recombinant proteins, we found that complete removal of the R2-C domain in RNF216 reduced total RNF216 self-ubiquitination (Figure 2B). This reduction was also found when expressing a FLAG-tagged version of RNF216 that lacked the R2-C region in HEK293 cells (Figure 2C).

**Figure 2:**
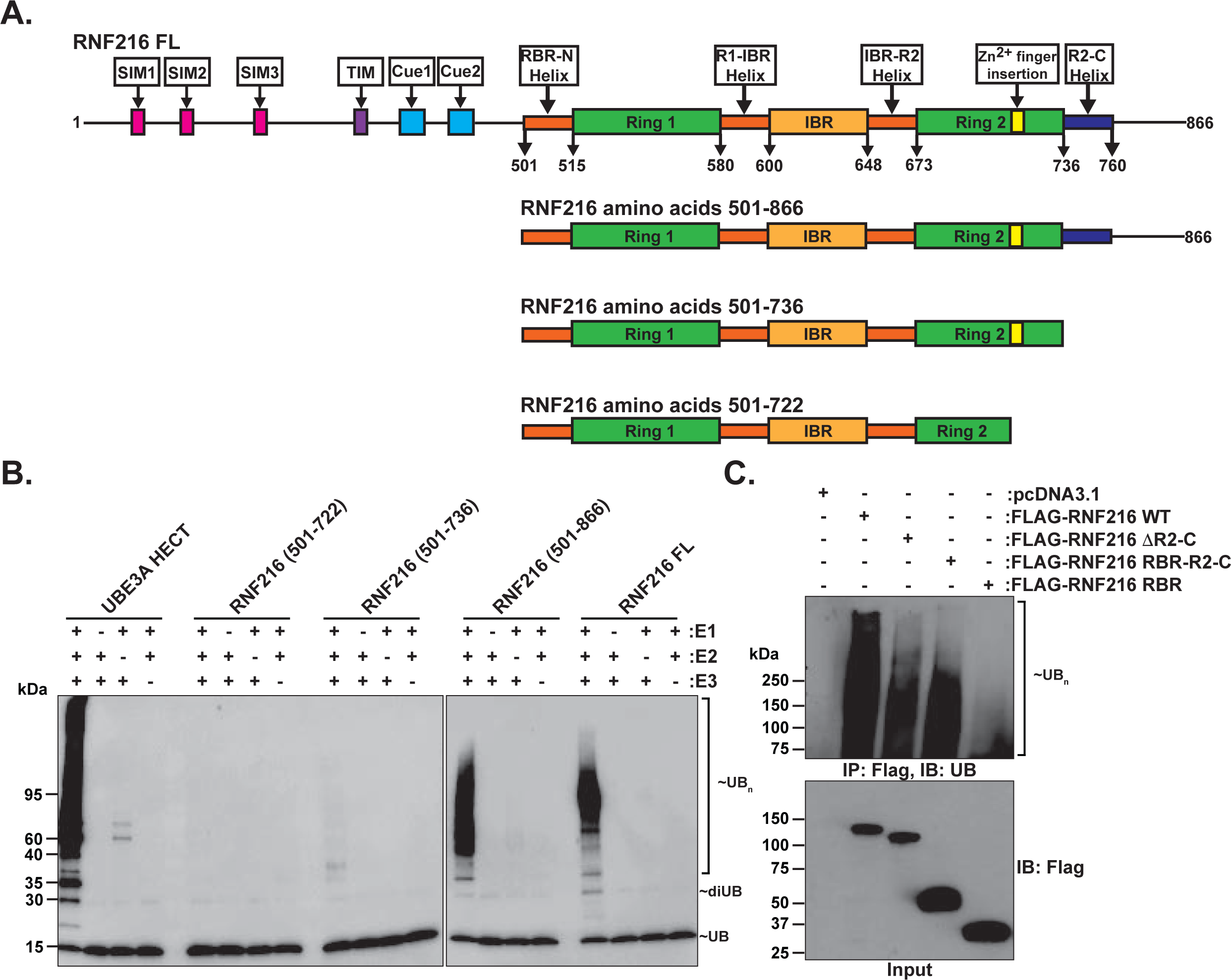
RNF216 R2-C domain is required for RNF216 self-ubiquitinating activity. (A) Domain structure of RNF216 and RNF216 truncation mutants created for in vitro studies. RNF216 contains a RBR (Ring 1, IBR, Ring2) domain with a Zn^2+^ finger insertion within Ring 2. The R2-C region with some homology to the LDD of HOIP is located between amino acids 736-760 and forms an α-helix when RNF216 is bound to a single ubiquitin (UB). (B) In vitro ubiquitination assay of RNF216 full-length (FL) and mutants. The HECT domain of UBE3A was used as a positive control. (C) FLAG-tagged versions of RNF216 were expressed in HEK293 cells. Cell lysates were immune-purified under denaturing conditions using an anti-FLAG antibody. Representative immunoblots probed with an anti-ubiquitin antibody. Note that removal of the R2-C domain decreased RNF216 autoubiquitination.

### RNF216 R2-C is required for RNF216 self-ubiquitination and ubiquitination of the RNF216 substrate Arc

Next, we examined the role of the R2-C domain in the assembly of ubiquitin chain types using ubiquitin linkage selective antibodies and ubiquitin mutants. Using the HECT domain of the E3 ligase UBE3A as a positive control for the assembly of K48-linked ubiquitin chains and a negative control for the assembly of K63-linked chains (22, 23), we found that RNF216 had no in vitro activity for assembling K48-ubiquitin linkages (Figure 3A, middle) but had strong activity for generating K63-linkages (Figure 3A, right). As expected, K63-linked ubiquitin activity was completely absent upon deletion of the entire R2-C domain. Mutation of the K63 site of ubiquitin to R to block polymeric ubiquitin assembly dramatically reduced RNF216 ubiquitination (Figure 3B, right), which was not as severe upon mutation of the K48 site of ubiquitin to R (Figure 3B, middle). However, once again, any activity was abolished when the complete R2-C domain was deleted. Taken together, these findings strongly indicate that the LDD-like domain of RNF216 dominantly functions to regulate global ubiquitination of RNF216 with a dominance toward forming K63-linked chains.

**Figure 3:**
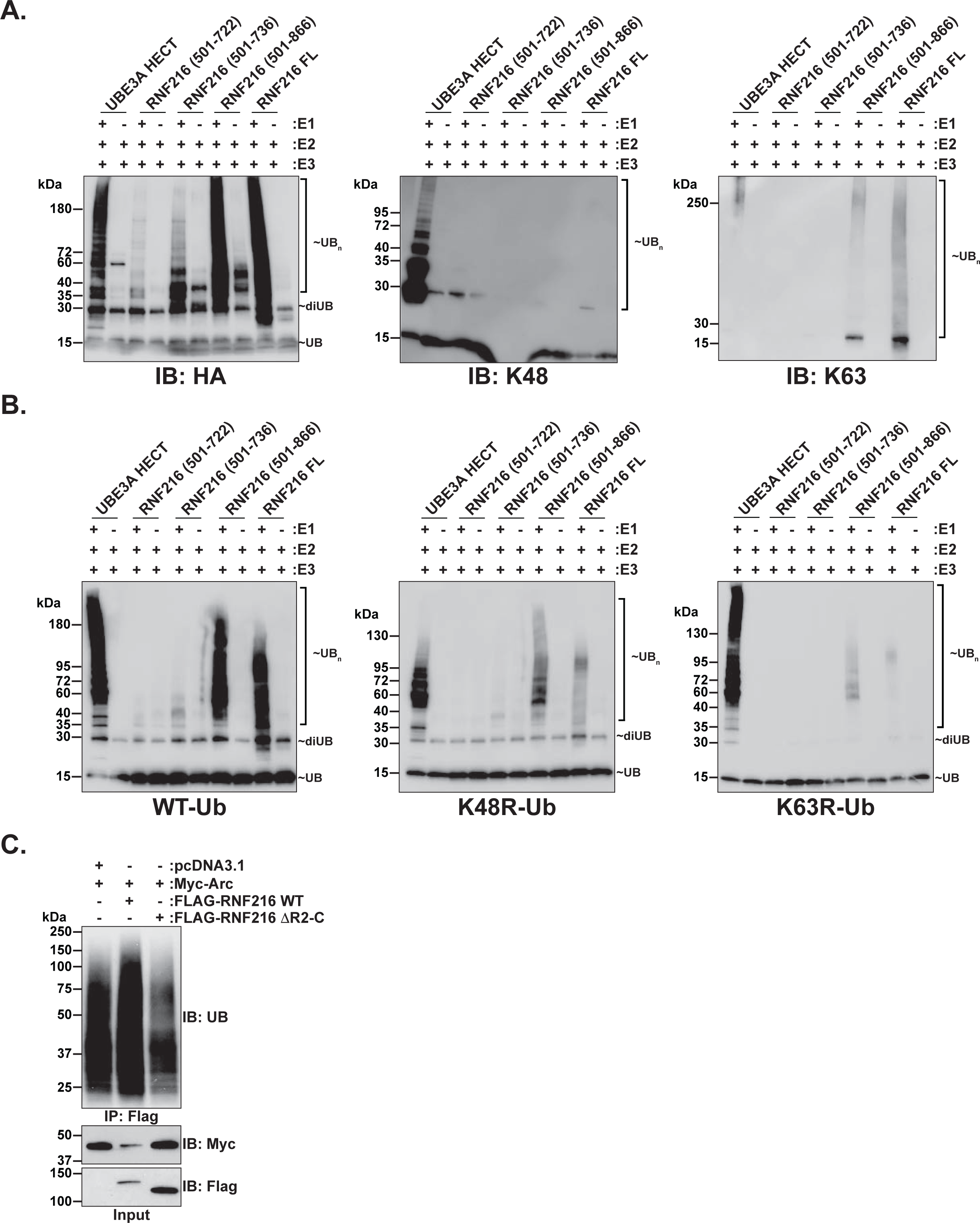
RNF216 R2-C domain reduces K63-linked self-assembly and is required for Arc ubiquitination. (A) In vitro ubiquitination assay of RNF216 full-length (FL) and mutants. Immunoblots from reactions were probed with anti-HA (left), anti-K48-specfic (middle), or anti-K63-specific (right) antibodies. Purified FL-RNF216 only had activity for K63-linkages, which was abolished upon deleting the R2-C domain. (B) In vitro ubiquitination assay of RNF216 FL and mutants using mutated ubiquitin in the reaction. Lysine residue K48 or K63 was mutated to arginine in ubiquitin to prevent the assembly of these specific chains. Immunoblots from reactions were probed with an anti-HA antibody. RNF216 activity was mostly affected in the K63R ubiquitin mutant demonstrating its dominance to assemble K63-UB linkages. (C) RNF216 and RNF216 lacking the R2-C domain was co-expressed with Myc-Arc in HEK293 cells. Arc was immune purified under denaturing conditions. Representative immunoblots probed with an anti-ubiquitin antibody. Note that removal of the R2-C domain reduced Arc ubiquitination.

An increase in the self-ubiquitination of E3s is typically an indicator of their catalytic activity (24). Therefore, we next tested the role of the R2-C domain on its ability to ubiquitinate known substrates. We previously showed that the immediate early gene Arc is a RNF216 substrate (19). To investigate the role of R2-C on Arc ubiquitination, we co-expressed RNF216 and R2-C domain truncation mutants with Myc-tagged Arc in HEK293 cells. As shown previously, addition of RNF216 led to a robust increase in the ubiquitination of Myc-Arc (19, 20). As expected, this enhancement was reduced upon deletion of the R2-C domain in RNF216 (Figure 3C) demonstrating that the R2-C domain of RNF216 also promotes the ubiquitination of a known substrate. Based on the published crystal structures (7, 21) and these findings, the R2-C domain may help coordinate Zn^2+^ within the Ring 2 domain and may help to facilitate the binding of UB to the IBR-R2 α-helix.

### The RNF216 R2-C domain alters the localization of RNF216 in primary hippocampal neurons

To determine how the R2-C domain of RNF216 may affect its expression in primary hippocampal neurons, we took advantage of a *Rnf216* conditionally floxed (*Rnf216^fl/fl^*) mouse line (25). To remove endogenous *Rnf216*, we transfected primary *Rnf216^fl/fl^*hippocampal neurons with CRE recombinase in combination with mutant versions of Flag-tagged RNF216. Previously, we found that RNF216 played an important role in the trafficking of receptors in neurons (25). A fraction of RNF216 was also found to colocalize with clathrin-rich regions in both dendrites and dendritic spines indicating that RNF216 is at hotspots for endocytosis (19). To determine if the R2-C domain altered RNF216 localization at endocytic hotspots, we co-transfected *Rnf216^fl/fl^* primary hippocampal neurons with a neuronal-specific CRE bicistronic tdTomato reporter (CamKIIα-CRE-P2A-tdTomato), Flag-tagged RNF216 WT and mutant plasmids and Clathrin-dsRed (Figure 4A). FLAG-RNF216 and CA (catalytically inactive) was broadly expressed in neurons with strong localization in the cell body and a punctated distribution in dendrites and dendritic spines (Figure 4B). However, the ΔR2-C was more enriched in the nucleus. All proteins had a punctate localization in neuronal dendrites and to dendritic spines. Together, these results show that deletion of the R2-C domain leads to an enrichment of RNF216 in the soma but does not grossly alter RNF216 localization in dendrites and dendritic spines.

**Figure 4:**
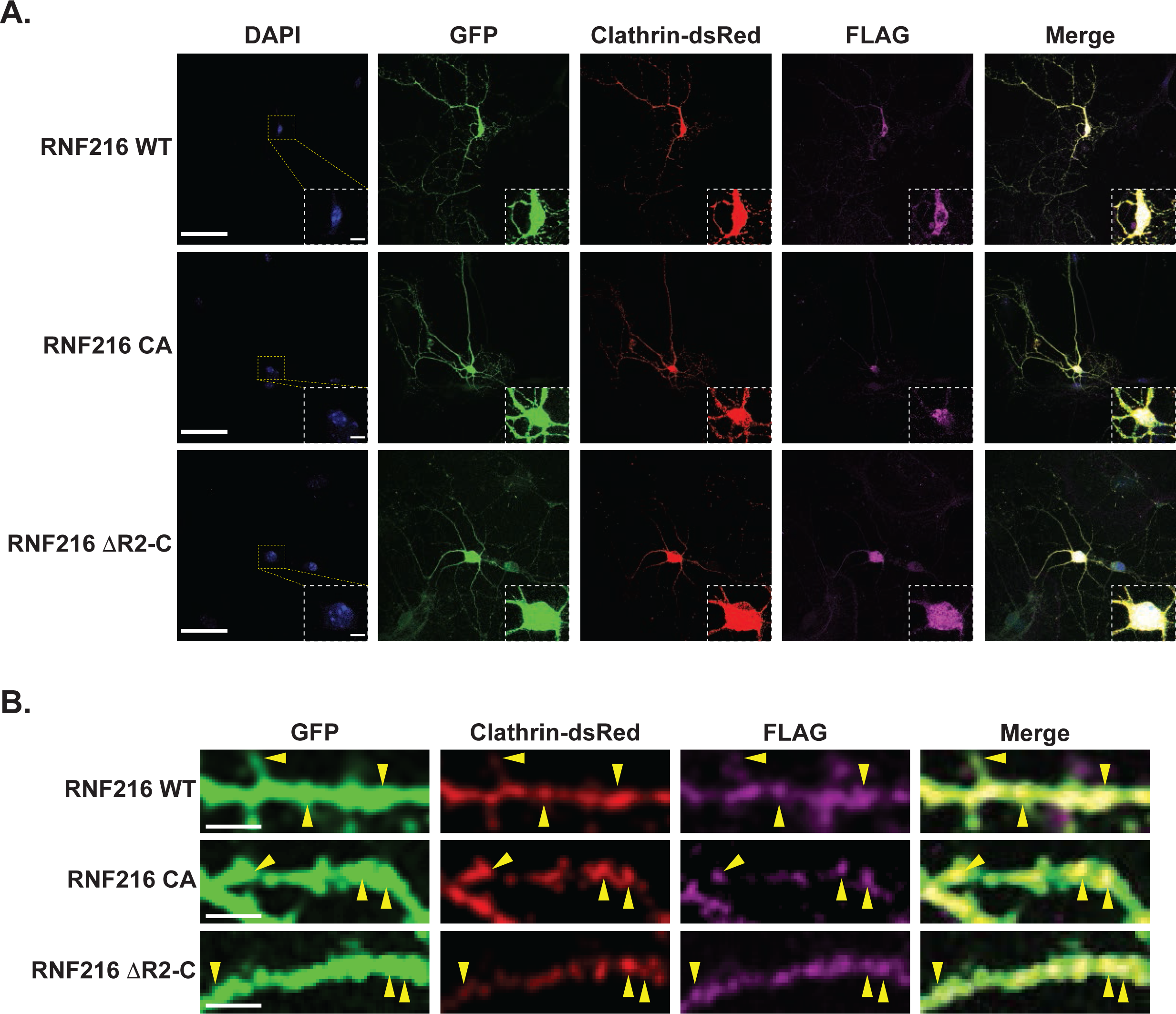
The RNF216 R2-C domain has greater nuclear localization in primary neurons but does not have altered localization in dendrites. (A) RNF216 localization in primary hippocampal neurons. *Rnf216^fl/fl^* primary hippocampal neurons were transfected at DIV13 with plasmids expressing CamKIIα-CRE-P2A-tdTomato (to delete endogenous RNF216 and to outline neuronal morphology) and FLAG-tagged versions of RNF216. Neurons were fixed and immunostained with anti-FLAG antibodies. *Inset*, Removal of the R2-C domain increased the expression of RNF216 in the nucleus. Scale bar, 50 μm, inset scale bar, 5 μm. (B) RNF216 localization in dendrites. Zoomed in images from A demonstrating localization of RNF216 in dendrites and with Clathrin-dsRed. Scale bar, 5 μm.

### The R2-C domain regulates dendritic complexity and dendritic spine morphology in primary hippocampal neurons

To determine a functional role for the RNF216 R2-C domain in neurons, we evaluated neuronal morphology upon deletion of the R2-C domain when expressed in *Rnf216^flfl^* CRE-transfected hippocampal neurons. Removal of the R2-C domain resulted in a reduction in the number of dendritic intersections as measured by Sholl analysis (Figures 5A-5B) whereas expression of the RNF216 catalytic inactive mutant (CA) was not significantly different from WT. The ΔR2-C mutant also had a significant increase in the total dendritic length compared to WT and CA expressing neurons (Figure 5C). These findings indicate that the R2-C domain uniquely regulates dendritic length and cell complexity.

**Figure 5:**
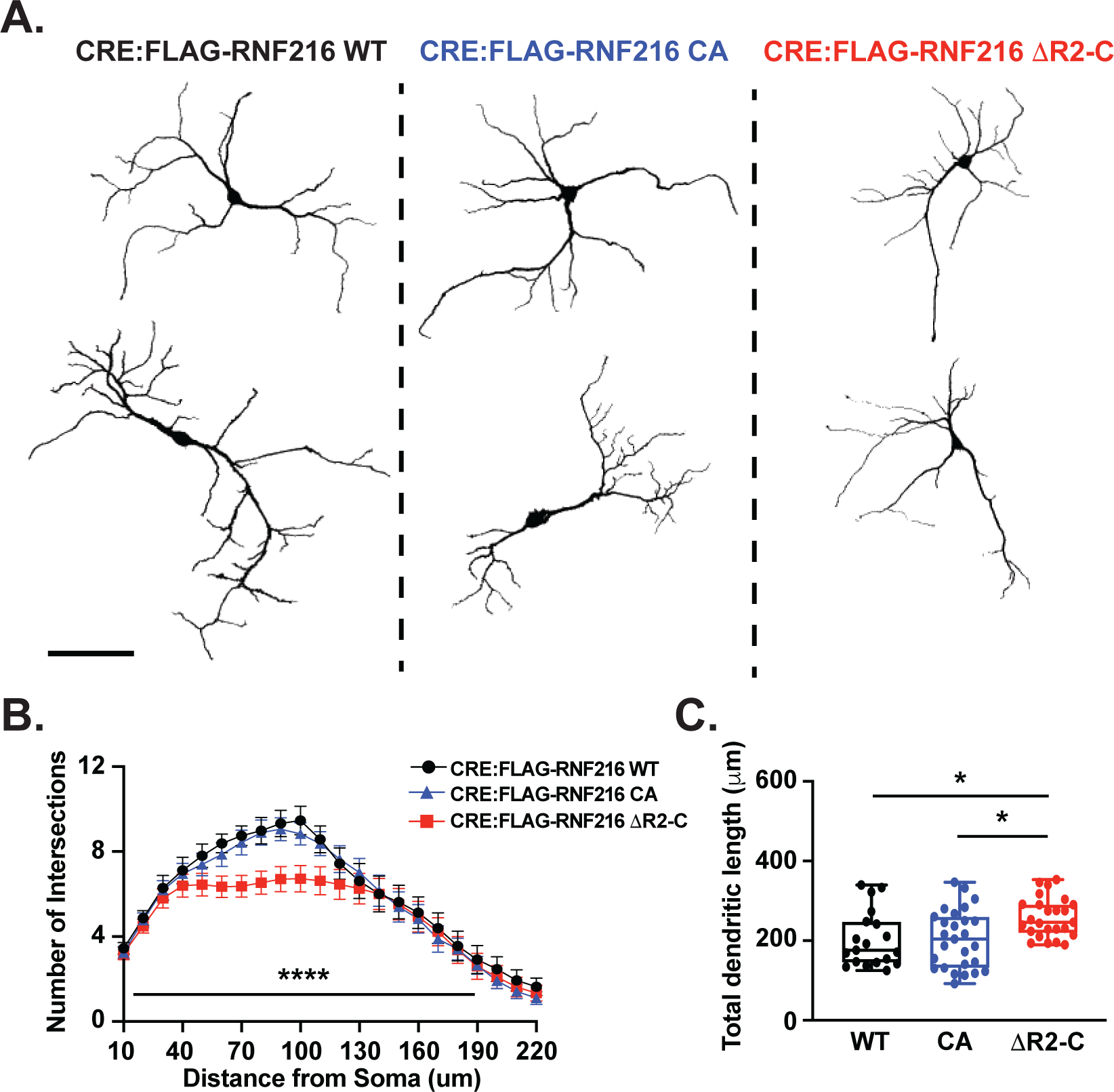
The R2-C domain selectively regulates dendritic morphology and size. (A) *Rnf216^fl/fl^* primary hippocampal neurons were transfected at DIV13 with plasmids expressing CamKIIα-CRE-P2A-tdTomato (to delete endogenous RNF216 and to outline neuronal morphology) and FLAG-tagged versions of RNF216. Neurons were fixed at DIV17, imaged and then outlined. Representative images of traced neurons from each experimental group. Scale bar, 50 μm. (B) Sholl analysis showing number of intersections for CRE:RNF216 WT (black), CRE:RNF216 CA (blue), and CRE:RNF216 ι1R2-C (red). Deletion of the R2-C domain significantly decreased the number of intersections. N = 19 – 27 per group, mixed-effects analysis, Condition F (1.973, 1517) = 15.20, ****p < 0.0001; Tukey’s multiple comparisons test of CRE:RNF216 WT vs. CRE:RNF216 ι1R2-C, ****p < 0.0001. (C) CRE:RNF216 ι1R2-C displays a significantly greater total dendritic length compared to CRE:RNF216 WT or CRE:RNF216 CA neurons. N = 19 – 27 per group, One-way ANOVA; Tukey’s multiple comparisons test CRE:RNF216 WT vs. CRE:RNF216 ι1R2-C, *p = 0.0197; CRE:RNF216 CA vs. CRE:RNF216 ι1R2-C, *p = 0.0101.

Previously, RNF216 was shown to regulate α-amino-3-hydroxy-5-methyl-4-isoxazolepropionic acid (AMPA) receptor abundance and regulate excitatory synaptic transmission at synapses (19), indicating that RNF216 may also regulate dendritic spine density or size. We measured the effects of the R2-C domain of RNF216 on dendritic spine densities and morphologies in hippocampal basal and apical dendrites. In basal dendrites, we did not observe a change in total spine density, spine type, or spine volume between each condition (Figure 6A-D). In contrast, while there were no significant differences in total spine density between each condition in apical dendrites (Figure 6E-F), removal of the R2-C domain displayed a significant higher density of stubby spines and a significant lower density of long thing spines compared to the RNF216 WT condition (Figure 6G). Dendritic spine volumes were not significantly different in any of the conditions (Figure 6H). Cumulatively, we conclude that the R2-C domain acts independently of the RBR catalytic domain to alter dendritic complexity and apical dendritic spine types.

**Figure 6:**
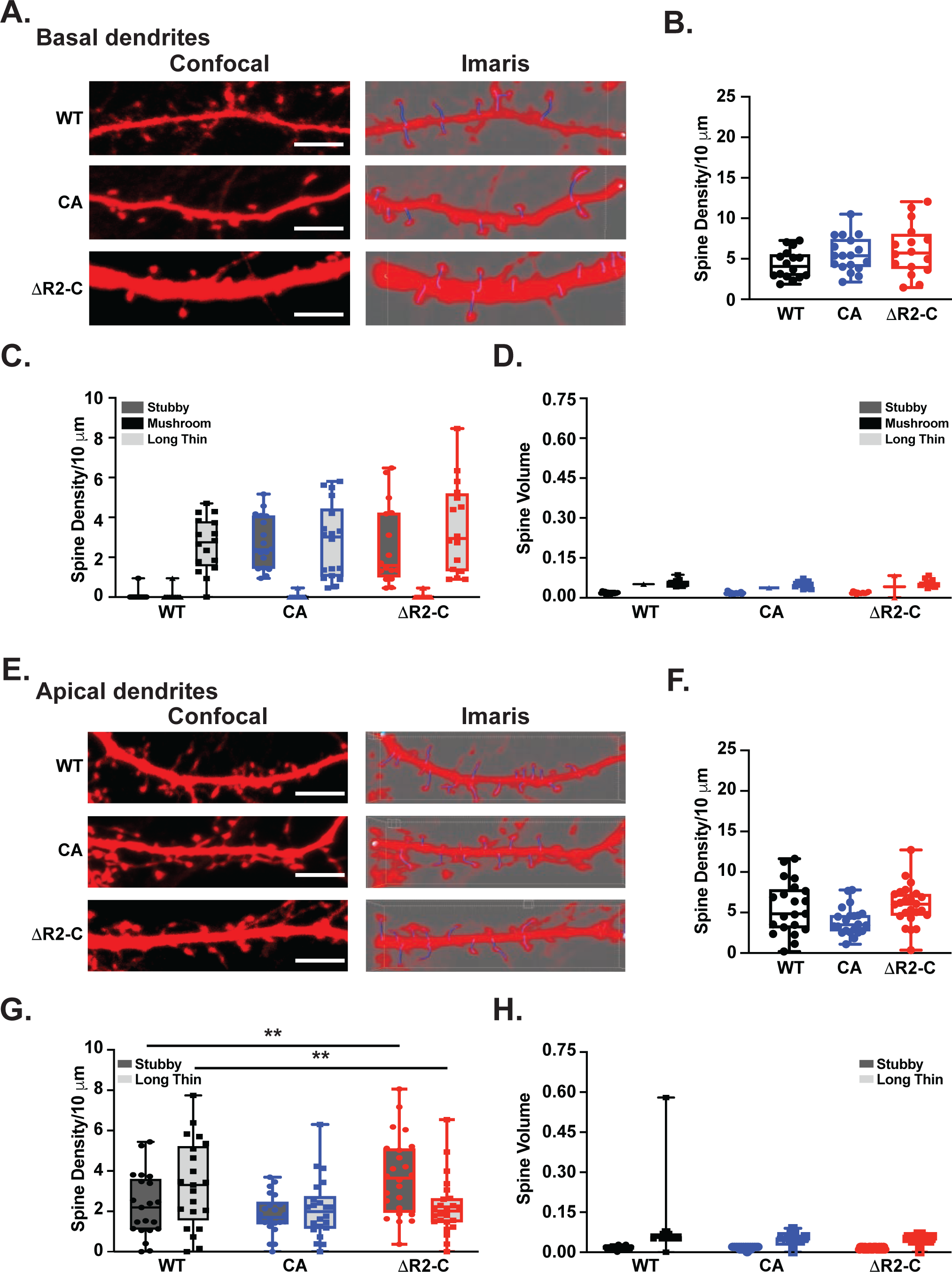
The R2-C domain does not alter dendritic spine type or densities in hippocampal neuron basal dendrites but does increase the density of stubby spines in apical dendrites. (A) *Rnf216^fl/fl^* primary hippocampal neurons were transfected at DIV13 with plasmids expressing CamKIIα-CRE-P2A-tdTomato (to delete endogenous RNF216 and to outline neuronal morphology) and FLAG-tagged versions of RNF216. Neurons were fixed at DIV17, imaged and then outlined. Representative images of dendritic segments of neurons from basal dendrites (left). Reconstructed neuron images from Imaris (right) showing long thin spines (blue) and stubby spines (pink). Scale bar, 5 μm. (B) No difference in total dendritic spine density between groups. N = 16 – 18 reconstructed dendritic segments per neuron per group. (C) No difference in the density of dendritic spine type per group. N = 16 – 18 reconstructed dendritic segments per neuron per group. (D) No difference in dendritic spine volumes between each group. N = 16 – 18 reconstructed dendritic segments per neuron per group. (E) *Rnf216^fl/fl^* primary hippocampal neurons were transfected as described in A. Scale bar, 5 μm. (F) No difference in total dendritic spine density between groups. N = 21 – 24 reconstructed dendritic segments per neuron per group. (C) Increased density of stubby spines in CRE:RNF216 ΔR2-C neurons. N = 21 – 24, One-way ANOVA; Tukey’s multiple comparisons test CRE:RNF216 WT vs. CRE:RNF216 ΔR2-C, **p = 0.0093. Decreased density of stubby spines in CRE:RNF216 ΔR2-C neurons. N = 21 – 24, One-way ANOVA; Tukey’s multiple comparisons test CRE:RNF216 WT vs. CRE:RNF216 ΔR2-C, **p = 0.0092. (D) No difference in dendritic spine volumes between each group. N = 21 – 24 reconstructed dendritic segments per neuron per group.

## Discussion

The ubiquitin pathway plays a major role in cellular signaling and biological processes that include neurodevelopment and neurotransmission (2, 26). E3 enzymes are major contributors to the diversity of biological functions for protein ubiquitination however how their domain structure supports the diversity and variety of ubiquitin chains is unclear. Here, we characterized a region within the RBR E3 RNF216 that shared homology to the LDD domain of the RBR E3 HOIP. While we expected this region to function in a similar manner to HOIP in assembling linear ubiquitin chains, we instead found that this domain was necessary to support the formation of all assembled RNF216 polyubiquitin chains. Recent evidence indicates that RNF216 has the capacity to assemble K11-, K48-, and K63-linked chains with a stronger preference for assembling K63-linkages (7–9). Using purified RNF216, we also found that it preferred to assemble K63-linkages in vitro as opposed to K48-linkages. These findings are in agreement with numerous studies that demonstrate a bias in RNF216 forming K63-ubiquitin linkages (7–9). Moreover, removal of the LDD-like domain of RNF216 abolished its ability to assemble any type of ubiquitin chain and was found to be essential to ubiquitinate its known substrate Arc (19). Together, these findings indicate the C-terminal region and its assembly has diverged from the LDD domain of HOIP and further validates HOIP as the only E3 that has capacity to assemble linear ubiquitin chains.

To establish a function of the R2-C domain in neural regulation, we characterized changes in its localization patterns in primary hippocampal neurons and determined its effects on neuron morphologies. We previously showed that RNF216 was broadly distributed in neurons in regions that included the pre– and postsynaptic regions, the neuronal soma, and in dendrites and dendritic spines. RNF216 also co-localized with clathrin in areas that were enriched at endocytic hotspots within spines called endocytic zones (19). Using TIRF microscopy, RNF216 also stably localized with clathrin at the cell surface (19) and further regulates the trafficking of receptors in dendrites, which was dependent on its RBR catalytic activity (CA) (25). Based on these findings, we set out to determine if the R2-C or the catalytic RBR region (CA) of RNF216 altered its localization. While removal of the R2-C and mutation of the RBR region (CA) did not result in changes in localization within dendrites and dendritic spines, there was an effect on the distribution of RNF216 in the nucleus where removal of the R2-C domain led to its nuclear accumulation.

Next, we determined if the R2-C domain altered the developmental profiles of neurons. Mutations in *RNF216* are found in the rare neurodegenerative disorder Gordon Holmes syndrome (GHS) (27). GHS individuals with RNF216 mutations exhibit hypogonadotropic hypogonadism, ataxia, dementia, and gray and white matter reductions. Mutations within the RBR domain and frameshift mutations that lie at the C-terminal region of RNF216 have been identified in GHS patients (27–31). Furthermore, RBR mutations have been shown to reduce the catalytic activity of RNF216 and its ability to ubiquitinate its known substrates (8, 32). However, frameshift mutations affecting the R2-C domain have not been studied. Here, we were surprised to find that the removal of the R2-C domain had differential effects on neuron morphologies compared to the catalytic RBR mutation (CA). Removal of the R2-C domain reduced dendritic complexity and increased overall dendritic length compared to WT while expression of the CA mutant was not significantly different from WT. Moreover, removal of the R2-C domain increased the density of stubby dendritic spines and reduced the number of long thin spines selectively in apical dendrites whereas the CA mutant was not significantly different from WT. While long thin spines are more dynamic and less stable, stubby spines are often more stable (33) indicating that the absence of the R2-C domain leads to premature maturation and stability of dendritic spines in apical dendrites. This accelerated development could confer resistance to synaptic plasticity that are necessary during critical periods of development. Unfortunately, we were not able to evaluate the formation of mushroom spines since our hippocampal neuron cultures at this age do not develop this spine type. However, it is worth noting that we were able to detect the formation of mushroom spines in basal dendrites lacking the R2-C domain (data not shown), which further supports the role of the R2-C domain in controlling dendritic spine development. Nevertheless, these findings suggest that differential mutations in GHS may lead to heterogenous phenotypes. Given that the R2-C domain increases RNF216 localization to the nucleus, we propose that removal of this domain may alter nuclear functions thus creating a gain-of-function phenotype, ultimately affecting neuronal development processes that include dendritic branching and apical dendritic spine maturation.

Based on the published crystal structures of RNF216 with ubiquitin (7, 21), the R2-C region forms an RBR-C-helix that lies in close proximity to the IBR-R2-helix that is bound to ubiquitin and is near a disordered region that lies adjacent to a Zn^2^ ion that is part of the RBR. The ability of RBR domains to coordinate with Zn^2+^ is an essential function for RBR E3 catalytic activity (34). Thus, the R2-C domain could serve as an allosteric modulator, stabilizing RBR coordination with ubiquitin while also stabilizing disordered RBR structure with Zn^2^. Interestingly, the RBR HHARI had similar homology to the RNF216 R2-C domain so this region may also function similarly to regulate HHARI ubiquitinating activity. Future directions would be to attempt to crystalize RNF216 with ubiquitin in the absence of the R2-C domain.

Cumulatively, our findings demonstrate that the R2-C domain of RNF216 has a unique role in regulating RNF216 ubiquitinating activity and functions to regulate dendritic morphology during development. Our work also highlights the possibility that mutations affecting different domains of RNF216 may have differing effects on neurodevelopmental profiles, creating a diversity of phenotypes in individuals with GHS.

## Acknowledgements

We would like to thank Dr. Dan Cox and Kevin Donaldson for Imaris resources and guidance. This work was funded by a NSF Career Award (2047700) and the Cleon C. Arrington Research Initiation Grant Program (RIG-93) to A.M.M.; the NINDS (grant R21NS116760) and a Molecular Basis of Disease Seed Grant to A.M.M. and J.Y. The work is also funded by NIH (R01GM104498) and NSF (2109051) to J.Y.

